# Immunologic Effect of Polysaccharides Extracted from *Sipunculus nudus* (SNP) on Hepatoma HepG2-bearing Mice

**DOI:** 10.1101/175190

**Authors:** Jie Su, Linlin Jiang, Jingna Wu, Zhiyu Liu, Yuping Wu

## Abstract

Since many studies have clarified the biological activity of polysaccharides, we investigated the effect of SNP which was the water-soluble polysaccharides extracted from *Sipunculus nudus* on Hepatoma HepG2-bearing Mice to verify the potential of SNP as an effective clinical agent for liver cancer therapy. SNP were administered at the doses of 50,100, and 200 mg/kg to HepG2-bearing mice to determine their antitumor effects. SNP had an inhibitory effect on the growth of HepG2 cells and enhanced the immunological effect on HepG2 tumor-bearing mice. SNP increased the expression of IL-2, IFN-γ, and TNF-α cytokines in serum, suggesting that SNP can strengthen the antitumor immune response. In addition, SNP increased ATF4, DDIT3, and IkBα expression and decreased CYR61, HSP90, and VEGF expression, all of which are proteins involved in antitumor activity and cell death/survival. our results suggested that SNP may be a novel antitumor agent.

**Summary statement:** SNP(polysaccharides extracted from *Sipunculus nudus*) mediates anti-tumor activity through influencing immunoregulation, and SNP can be explored as a promising candidate for future anticancer drug.

## Introduction

Hepatitis B (HBV)-associated hepatoma is one of the leading causes of cancer death in Asia(Sun *et al*, 2011). Currently, the treatment of hepatoma includes surgery and intervention chemotherapy; however, these measures often have a variety of side effects (Paraskevi *et al*, 2006). Therefore, identification of non-invasive treatments is in great interest for hepatoma therapy that may increase the survival quality of cancer patients.

SNP, the main ingredient in *Sipunculus nudus* extract, is a monosaccharide that mainly comprises L-rhamnose, L-arabinose, D-ribose, D-glucose and D-galactose(Liu *et al*, 2011). *In vivo* and *in vitro* studies have indicated that SNP intake may enhance immune function (Cui *et al*, 2014). We previously demonstrated the pro-apoptotic activity and anti-HBV virus activity of SNP on HepG2.2.15 cells (Su *et al*, 2016). In the present study, we show that SNP can stimulate immunological function to mediate antitumor activity. Further development and exploration of SNP may identify SNP as an effective antitumor agent for clinical liver cancer therapy.

## Materials and methods

### Isolation and purification

Referring to previous reports (Su *et al*, 2016), *S.nudus* (diameter 6 ± 2 mm, length 10 ± 2 cm) was collected from Xiamen market, China. Fresh peanut worms were washed, oven dried, and crushed into powder. The powder was hydrolyzed in trypsin at 50 °C for 5 hours, and the insoluble material was removed by centrifugation. The supernatant liquor was deproteinized 4 times using the Sevage method(Staub,1965; ^a^ Zhang *et al*, 2011). Next, 4 volumes of cold ethanol was added to precipitate the material after standing at 4 °C overnight (^b^ Zhang CX *et al*, 2011). The resulting precipitate was then centrifuged (3000 g × 10 min). After washing with ethanol 3 times, the precipitate was freeze-dried *in vacuo* and ground into a powder to produce a crude product. The crude product was subjected to a DEAE-Sepharose anion exchange column (3.0 cm × 40 cm), eluting at 0.5 ml/min successively with 0.0175 M pH 6.7 phosphate buffer solutions. Each fraction was collected with 2 ml of elute. The main elution fraction containing the carbohydrates was concentrated, dialyzed, and lyophilized.

### Mice assay

Male BALB/c mice (6-8 weeks old; 18˜22g) were obtained from Shanghai Slack Laboratory Animal Co. Ltd. All mice were housed under conditions of 20-25□ room temperature, 30%-70% relative humidity, and 12-hour light/dark cycle. Animals were fed with the standard mice food and provided tap water *ad libitum*.

### Cell culture

The HepG2 cell line was obtained from ShangHai MeiXuan Biological Technology Co. Ltd. and cultured in DMEM medium with 10% fetal bovine serum (FBS). Cells were incubated at 37□ in 5% CO_2_ humidified air.

### Experimental models

A total of 100 mice were used in this study and divided into two parts: treatment and prevention. For the treatment group, 50 mice received a subcutaneous inoculation of HepG2 cells (2×10^6^ cells per mouse). A week after tumor cell inoculation, we screened 30 mice for millet size tumors and used as the experimental animals. The experimental animals were randomly divided into five groups of 6 mice each----vehicle control (saline), astragalus polysaccharides (APS; 100 mg/kg), and three SNP groups (50,100, and 200 mg/kg).

For the prevention study, the other 50 mice were randomly divided into four groups---preconditioning vehicle and three SNP groups (50,100, and 200 mg/kg). The mice were initially administered drugs according to respective treatment groups by intragastric administration for 1 month. Next, the animals received a subcutaneous inoculation of HepG2 cells (2×10^6^ cells per mouse). A week after injection, 30 mice were screened for millet size tumors, divided into 6 mice/group and used as the experimental models.

### Antitumor assays

Drugs were given per group once daily intragastrically for 16 consecutive days after tumor screening. The vehicle group consisted of normal saline intake mice (0.2 ml each mice).

All animals were humanely killed and tumors, thymuses, and spleens were dissected and weighed at the end of the assay. Body weights, tumor weights, thymuses weights and spleens weights were recorded. The tumor inhibition rate and organ index were calculated using the following equation:

Tumor inhibition rate = (mean tumor weight of vehicle group – mean tumor weight of treated group)/mean tumor weight of vehicle group×100%

Organ Index=organ weight/body weight×100%

### Histology

At the end of the antitumor assay, tumors of all mice were collected for histological examination. These tissues were fixed in 4% paraformaldehyde solution, embedded in paraffin, and serially cut for hematoxylin and eosin (H&E) staining. Samples were imaged under a light microscope for morphological observation.

### Cytokines assay

At the end of the antitumor assay, blood samples were obtained from the sinus by extracting the eyes of the mice. After 2h’ incubation, blood serum was separated by centrifugation at 3000 rpm, 10min. Cytokines including interleukin (IL) 2, interferon (IFN) γ, and tumor necrosis factor (TNF) α were determined by enzyme linked immunosorbent assay (ELISA).

### Western Bloting

In the Western blot analysis, 50mg of each tumor was harvested, and these samples were added in cell lysis buffer. After trituration and centrifugation, the concentration of protein per group was determined using a BCA protein assay kit according to the manufacturer’s protocol. Then 20μg of total protein was used for SDS-PAGE, proteins were transferred to membranes, and blocked with 5% skim milk. ATF4, DDIT3, IkBa, CYR61, HSP90 and VEGF primary antibodies were added separately and the sections were incubated at 4°C overnight. Subsequently, membranes were washed in PBS and secondary antibodies were added and incubated correspondingly. After washing with PBS, the membranes were stained with ECL, and observed on an automatic electrophoresis gel imaging analyzer. Protein levels of above mentioned interest protein were analyzed by comparing with an internal reference.

### Statistical analysis

Data was expressed as means ± standard deviation (S.D.). Student’s *t* test was used to analyze the differences between the control and test groups. Differences were considered to be statistically significant at *P* < 0.05 and very significant at *P* < 0.01.

## Results

### Isolation and Purification

Based on previous reports, polysaccharides were obtained after fractionation on a DEAE-Sepharose anion exchange column from *S.nudus.* SNP comprised Ara 10.7%, Rha 12.6%, Gal 16.4%, Glu 31.3%, Xyl 18.2%, and Man 10.8 %(Su *et al*, 2016).

### The inhibitory effect of SNP on the growth of HepG2 cells and its immunological enhanced effect on HepG2 tumor-bearing mice

In order to determine the suppression effect on HepG2 tumor, drugs were administered once daily for 16 consecutive days after screening tumor. There were no significant differences in the body weight between groups. Compared to the vehicle group, the tumor weight of APS treatment group, two SNP treatment groups (100mg/kg, SNP 200mg/kg), all SNP prevention groups were significantly lower (*P* < 0.01). The tumor weight of the SNP 200mg/kg treatment group, and preconditioning SNP 200mg/kg prevention group were similar with the APS treatment group. The tumor inhibition rates by weight represented a similar trend and were as follows: APS group 54.67%, SNP 200mg/kg group 48.52%, and preconditioning SNP 200mg/kg group 59.30% (Figure. 1A).

**Figure 1.**
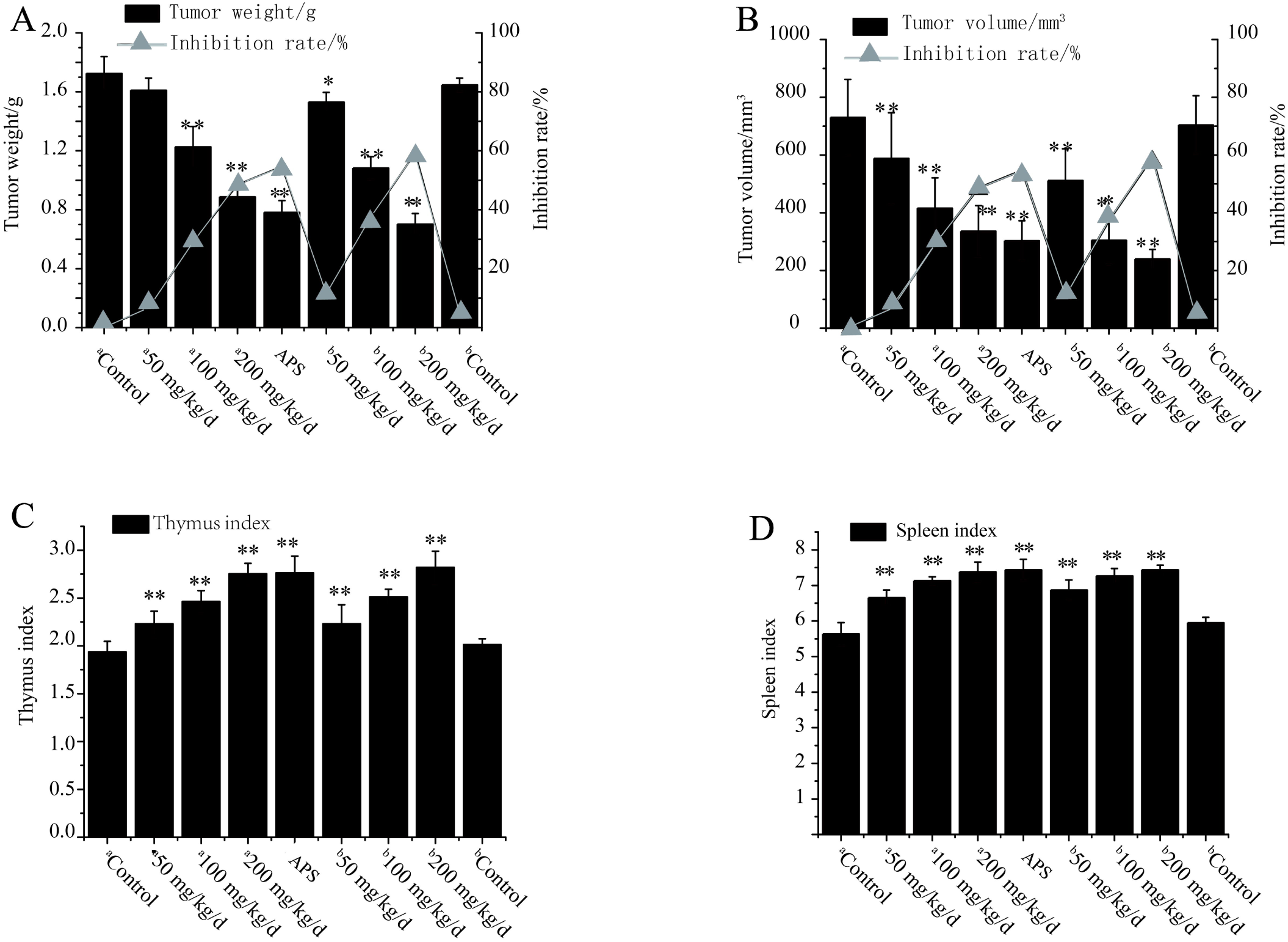
The inhibitory effect of SNPs on the growth of HepG2 cells and its immunological enhanced effect on HepG2 tumor bearing mice. Data were presented as mean ± SD (n=6). P value□<□0.05 compared to Vehicle group was marked *. P value□<□0.01 compared to Vehicle group was marked **. Not significant compared to Vehicle group was not marked.

Compared to the vehicle group, the tumor volume of APS treatment group, the three SNP treatment groups and preconditioning SNP 100mg/kg, SNP 200mg/kg prevention groups were significantly smaller (all *P* < 0.01), while the decrease in the preconditioning SNP 50mg/kg prevention group was also significant at *P*< 0.05. The tumor volume of SNP 200mg/kg treatment group, and preconditioning SNP 100mg/kg prevention group were similar with the APS treatment group, with a greater decline seen in the preconditioning SNP 200mg/kg prevention group. The tumor inhibition rates by volume were as follows: APS group 58.43%, SNP 200mg/kg group 53.99%, and preconditioning SNP 100mg/kg group 58.32%, preconditioning SNP 200mg/kg group 67.21%. These investigations indicated that SNP had an inhibitory effect on HepG2 tumor in mice (Figure. 1B).

Compared to the vehicle group, the thymus and spleen index values of APS treatment group, three SNP treatment groups, and the three preconditioning SNP prevention groups were significantly increased (*P* < 0.01). The thymus and spleen index values of SNP 200mg/kg treatment group and preconditioning SNP 200mg/kg prevention group were similar to the APS treatment group (Figure.1C, Figure.1D). These results suggested that apart from inhibitory effect, SNP also had an immunological enhanced effect on HepG2 tumor-bearing mice.

### Morphological Observation

The microscopic observation of the HepG2 tumor-bearing mice of vehicle treatment group and preconditioning vehicle prevention group both showed that the tumor cells were diffusely distributed, and arranged densely with different sizes. As anticipated, tumor cells of APS treatment group were restrained, arranged loosely and some tumor cells were swollen, with visible degeneration or even necrosis. In all six SNP (treatment and prevention) groups, we observed the similar histopathological changes with the APS treatment group (Figure. 2). These results indicated that SNP inhibited the proliferation of HepG2 tumor cells.

**Figure 2.**
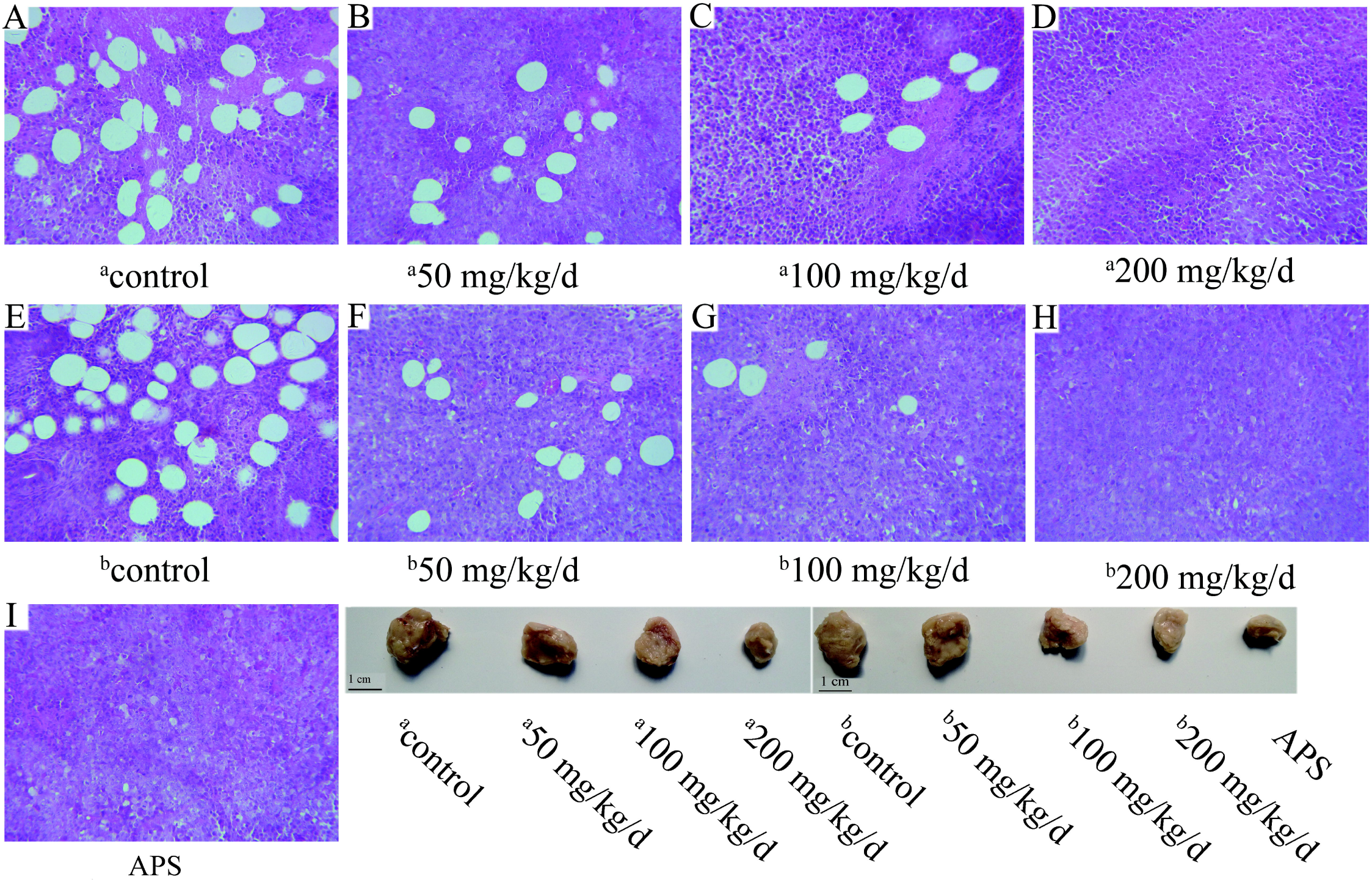
Morphological observation of tumor cells from HepG2 tumor bearing mice (HE stain). A, Vehicle group. B, SNPs 50 mg/kg group. C, SNPs 100 mg/kg group. D, SNPs 200 mg/kg group. E, Preconditioning vehicle group. F, Preconditioning SNPs 50 mg/kg group. G, Preconditioning SNPs 100 mg/kg group. H, Preconditioning SNPs 200 mg/kg group. I, APS group.

### The upregulated effect of SNP on Cytokines

The effect of SNP in IL-2, IFN-γ and TNF-α cytokines is shown in Figure 3. Compared to vehicle group, IL-2 expression in the APS treatment group and all six SNP groups was significantly upregulated (P < 0.01)(Figure. 3A). IFN-γ and TNF-α was significantly upregulated in the APS treatment groups, the three SNP treatment groups and two preconditioning SNP prevention g groups (100 mg/kg and 200 mg/kg) (all P < 0.01), while the increased expression in the preconditioning SNP 50 mg/kg prevention group was significant at P< 0.05(Figure. 3B, Figure. 3C). The expression of the three cytokines in the SNP 200 mg/kg treatment group and preconditioning SNP 200 mg/kg prevention group were similar with the APS treatment group. These investigations indicated that SNP had an upregulated effect on IL-2, IFN-γ and TNF-α cytokines in HepG2 tumors in mice.

**Figure 3.**
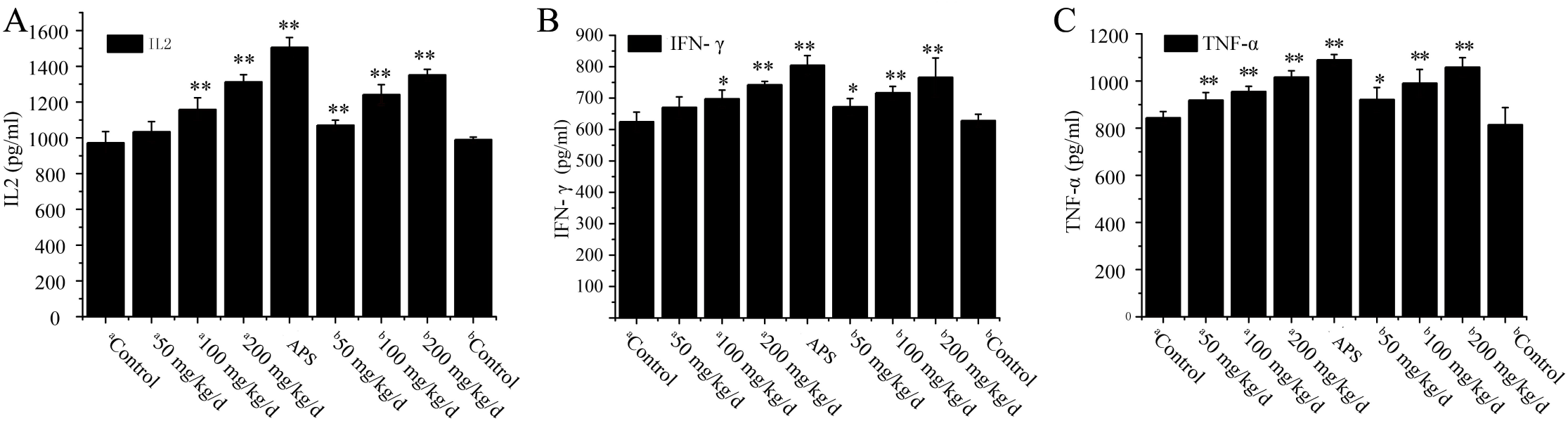
The up regulated effect of SNPs on Cytokines. Data were presented as mean ± SD (n=6). P value□<□0.05 compared to Vehicle group was marked *. P value□<□0.01 compared to Vehicle group was marked **. Not significant compared to Vehicle group was not marked.

### SNP Regulated the Protein Expression of ATF4, DDIT3, IkBa, CYR61, HSP90 and VEGF

The protein levels of ATF4, DDIT3, IkBa, CYR61, HSP90 and VEGF proteins are presented in Figure 4. The results showed that the protein levels of ATF4, DDIT3, and IkBa increased in the APS treatment group and all six SNP (treatment and prevention) groups compared to vehicle group. The protein levels of ATF4, DDIT3, IkBa in SNP 100 mg/kg treatment group were similar with the APS group, while the expression of these proteins were increased in both the SNP 200 mg/kg treatment group and the three preconditioning SNP prevention groups (Figure. 4A, Figure. 4C, Figure. 4E).

**Figure 4.**
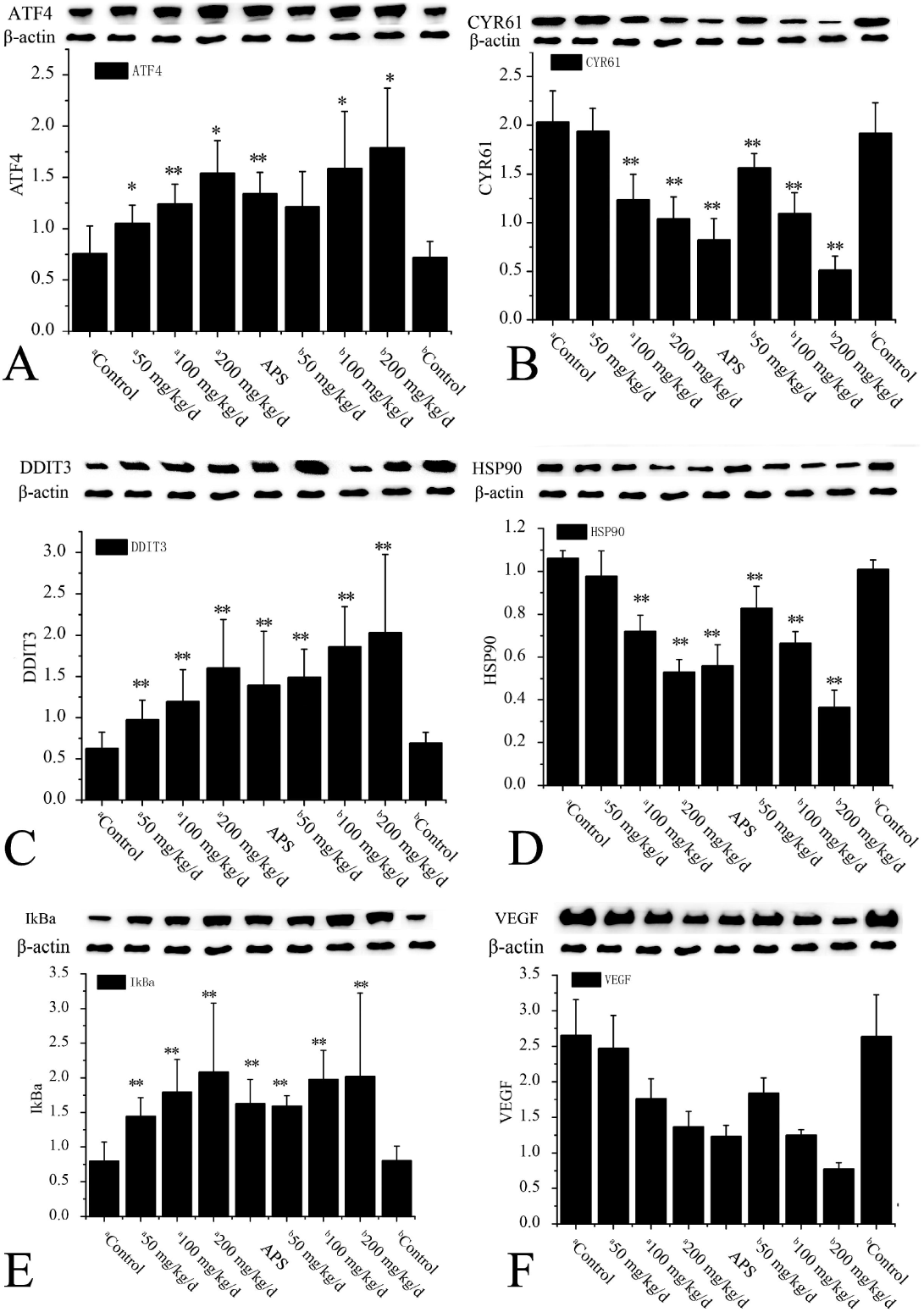
The regulated effect of SNPs in HepG2 tumor bearing mice using Western Blot assay. Data were presented as mean ± SD (n=6). P value□<□0.05 compared to Vehicle group was marked *. P value□<□0.01 compared to Vehicle group was marked **. Not significant compared to Vehicle group was not marked.

The protein levels of CYR61, HSP90, and VEGF in the APS treatment group, two SNP treatment groups (100 mg/kg 200 mg/kg) and the three preconditioning SNP prevention groups compared to the vehicle group. No differences were seen in the SNP 50 mg/kg treatment group compared to the vehicle group. The protein levels of CYR61, HSP90, and VEGF in SNP 200 mg/kg treatment group, and preconditioning SNP 100 mg/kg prevention group were similar with the APS treatment group, while the levels were significantly higher in the preconditioning SNP 200 mg/kg prevention group (Figure. 4B, Figure. 4D, Figure. 4F).

## Discussion

In our previous study, we demonstrated that the water-soluble polysaccharides (SNP) possessed anti-tumor activity, anti-virus activity (Su *et al*, 2016). Studies from other research groups have also shown the different potential immunologic effect of SNP (^a^Zhang *et al*, 2016; ^b^Zhang etal, 2016). In the present study, we isolated and purified SNP and explored their immunologic effect on hepatoma HepG2-bearing mice.

HepG2 cells have been used as a cellular tool to identify therapies for hepatitis B virus and hepatoma(Huang *et al*, 2016;Paramita *et al*, 2014). Here we established the hepatoma HepG2-bearing mice model to discern the immunologic effect of SNP. A recent study demonstrated that APS has therapeutic effect on hepatocellular carcinoma H22-bearing mice(Lai *et al*, 2017), thus, in this study we chose APS as the positive control to determine the significance of our findings.

First, we investigated the suppression effect of SNP on HepG2 tumor. Drugs were administered to each respective group once daily for 16 consecutive days after screening tumor. Similar to the APS treatment group, the tumor weight and tumor volume of two SNP treatment groups (100 mg/kg and 200 mg/kg) and the three preconditioning SNP prevention was significantly different from the vehicle group. These changes suggested that SNP could suppress the growth of HepG2 tumor in mice, and SNP treatment or prevention at 200 mg/kg had the same extent effect with APS group.

In a study by Zhao *et al.,* the thymus and spleen indexes were believed to reflect the immunological function of the organism (Zhao *et al,* 2013); therefore, we collected the data of thymus and spleen indexes to determine whether the impact of SNP tumor suppression effect was related to the immunomodulatory activity effect. In the APS treatment group and all SNP groups, the thymus and spleen index values were significantly increased compared to the vehicle group. Therefore, this data suggests that SNP could inhibit HepG2 tumor through the immunological enhanced effect, and 200 mg/kg SNP treatment or prevention had the same extent immunological enhanced effect as the APS group.

In the immunomodulatory activity of organisms or tumor development, cytokines and molecular pathways play pivotal roles, of which cytokines are the essential immunological response components to implement the antitumor role(Govaere *et al,* 2015; Jiang *et al,* 2015; Sun *et al,* 2014). It is believed that IL-2 is an important immunomodulatory cytokine involved in cells which reflect the cellular immunity; thus, increased IL-2 levels suggests enhanced cellular immunity in cancer patients(Zhao *et al*, 2013; Sun *et al*, 2014; Jin *et al*, 2012). In this study, IL-2 in all six SNP groups and the APS treatment group was significantly upregulated; this indicated that SNP could upregulate immune function. IFN-γ is a critical cytokine with both negative and positive activity under various circumstances (Chen *et al*, 2015). TNF-α is a pleiotropic cytokine that was originally described as anti-tumorigenic(Carswell *et al*,1975; Ichinose *et al*, 1988). The results in this study showed that IFN-γ and TNF-α in the APS treatment group, the three SNP treatment groups, and two doses in the preconditioning SNP prevention groups (100 mg/kg and 200 mg/kg) were significantly upregulated, indicating that SNP can strengthen the immune response by promoting the expression of IFN-γ and TNF-α.

Masuda *et al*. determined that TNF-α could induce the PERK-elF2α-ATF4-CHOP axis in the ER stress response(Masuda *et al*, 2013). Meanwhile, TNF-α inhibits activity of PPARγ through IkB*a-*mediated signaling in the pathogenesis of inflammation and cancer cachexia(Ye *et al*, 2008). ATF4 could promote transcription of many stress-related genes and one of its targets, DDIT3, can promote cell death(Tabas *et al*, 2011; Huggins *et al*, 2016). DDIT3 is known as a key regulator of cell stress response, and its target genes are involved in cell proliferation, apoptosis/survival, and cancer (Jauhiainen *et al*, 2012). Herein, we found that the gene expression level of ATF4, DDIT3 and IkBα in all six SNP groups was changed (data not shown) by the high-throughput sequencing assay; therefore, we speculated that SNP might be involved in two immunomodulatory signaling pathways to inhibit HepG2 tumor progression: (i) TNF-α-ATF4-DDIT3, and (ii) TNF-α-IkBa-PPARγ. Western blotting results in this study showed that the protein levels of ATF4, DDIT3 (also known as CHOP), and IkBα increased in all six SNP groups and APS treatment group compared to the vehicle group. This data confirmed that SNP can increase immune activity through regulating TNF-α, ATF4, DDIT3, and IkBα to inhibit HepG2 tumor progression.

Moreover, we found that the gene expression level of CYR61, HSP90, and VEGF decreased in all six SNP groups by high-throughput sequencing assay. CYR61 is related to cell survival, proliferation, and differentiation, and is over expressed in human tumors (Zhu *et al*, 2016). Hsp90 has been used as a target for anticancer therapeutics since 1994(Alarcon *et al*, 2012). In addition, VEGF target therapy is an important treatment for many human malignancies (Yamagishi *et al*, 2013). It is recognized that the over expression of these three proteins are induced in human tumors. In this study, we further determined the expression of these three protein levels to explore the potential of SNP as a promising candidate for future anticancer therapy. The results showed that the protein levels of CYR61, HSP90, and VEGF were decreased in the APS treatment group, two SNP treatment groups (100 mg/kg and 200 mg/kg), and the three preconditioning SNP prevention groups as compared to vehicle group. Therefore, we conclude that SNP plays an immunoregulatory effect in HepG2 tumor-bearing mice through inhibiting the expression of CYR61, HSP90, and VEGF.

It is noteworthy that the 200 mg/kg SNP treatment and prevention groups had the same extent of immunological enhanced effect as the APS treatment group as determined by the thymus and spleen indexes and cytokines assay. While the immunomodulatory effect of ATF4, DDIT3, IkBα, CYR61, HSP90, and VEGF of SNP 100 mg/kg treatment group were similar with the APS group, the levels in the SNP 200 mg/kg treatment group and the three preconditioning SNP prevention groups were increased. This inconsistent tendency suggests the following hypothesis: 1) the higher the dose, the more obvious the effect. However, in our study the highest dose of SNP was 200 mg/kg; therefore, a future study will be necessary to explore whether higher doses can exhibit higher efficacy without increasing toxicity.2) The regulation of ATF4, DDIT3, IkBa, CYR61, HSP90, and VEGF was significantly different between SNP treatment and prevention groups, indicating that vaccinating healthy mice with SNP can inhibit HepG2 tumor development to some extent; but the difference was only seen at the protein level, indicating that under our study conditions, SNP merely influenced the molecular level. Future studies should be done to determine how to magnify this effect to the whole organism.

## Conclusion

In this study, we demonstrated that SNP had immunologic effect on hepatoma HepG2-bearing mice. Our findings show that the SNP dose of 50-200 mg/kg inhibited the proliferation of HepG2 cells and increased the thymus and spleen indexes, also upregulated the IL-2, IFN-γ, and TNF-α cytokines in serum. Besides, SNP increased ATF4, DDIT3, and IkBα expression and decreased CYR61, HSP90 and VEGF expression, all of which are involved in anti-tumor activity and cell death/survival. In conclusion, our findings suggest that SNP mediates anti-tumor activity through influencing immunoregulation, and SNP can be explored as a promising candidate for future anticancer drug.

## Acknowledgements

We thank members of Key Laboratory of Cultivation and High-value Utilization of Marine Organisms in Fujian Province for assistance collecting *Sipunculus nudus*. We thank Shanghai Jiao Tong university animal research center for their animal experiment service. We thank Accdon (www.accdon.com) for its linguistic assistance during the preparation of this manuscript.

## Author contributions

All authors designed the study, were involved in data collection, and analyzed the data. Jie Su wrote the manuscript.

